# SkinnerTrax: high-throughput behavior-dependent optogenetic stimulation of *Drosophila*

**DOI:** 10.1101/080614

**Authors:** Ulrich Stern, Chung-Hui Yang

## Abstract

While red-shifted channelrhodopsin has been shown to be highly effective in activating CNS neurons in freely moving *Drosophila*, there were no existing high-throughput tools for closed-loop, behavior-dependent optogenetic stimulation of *Drosophila*. Here, we present SkinnerTrax to fill this void. SkinnerTrax stimulates individual flies promptly in response to their being at specific positions or performing specific actions. Importantly, SkinnerTrax was designed for and achieves significant throughput with simple and inexpensive components.

## Main text

Artificially stimulating or inhibiting neurons is an important tool for unraveling the neural mechanisms that control animal behavior. A widely used method of stimulation is optogenetics, where channelrhodopsin is expressed in the neurons of interest, allowing them to be activated with high temporal resolution using light pulses. In *Drosophila*, targeted expression of channelrhodopsin is made easy by techniques like the Gal4-UAS system, and no surgery is needed to enable light delivery when using red-shifted channelrhodopsin^1, 2^ as red light penetrates *Drosophila* sufficiently well. Several recent reports^3-6^ have demonstrated the effectiveness of this approach in assessing whether activating specific neurons is sufficient to drive specific behaviors. A more sophisticated application of optogenetics, however, is to stimulate neurons of interest with a closed-loop system in real-time response to the fly’s positions and actions. Such application is critical for elucidating the roles of specific neurons in several important behavioral tasks such as operant learning.

Existing systems that allow such behavior-dependent stimulation have not been designed for high throughput. Two recent advanced systems, FlyMAD^7^ and ALTOMS^8, 9^, target a laser to flies in real-time using mirror galvanometers, which in principle allows sharing the laser between several flies, as suggested in the discussion section of Bath et al.^7^ But the scalability of sharing the laser is limited (**Supplementary Note 1**), and laser-targeting systems require expensive components and complex setups. Thus, for a typical lab laser-targeting systems likely make achieving high throughput prohibitively difficult.

Here, we present SkinnerTrax, a system for high-throughput behavior-dependent optogenetic stimulation of *Drosophila*. Importantly, SkinnerTrax was designed for and achieves significant throughput with simple and inexpensive components. SkinnerTrax can deliver red light of different intensities and durations to individual flies promptly in response to their being at specific positions or performing specific actions. Instead of targeting a laser onto a fly that is to be stimulated, SkinnerTrax uses regular LEDs to illuminate the whole arena for such fly, greatly simplifying light delivery. For our setup, SkinnerTrax could handle 32 arenas (flies) simultaneously utilizing only 9% CPU of an Intel i7 machine and using a simple multi-channel LED controller. To scale up further, the number of arenas per machine or the numbers of machines and LED controllers can be increased easily.

Conceptually, SkinnerTrax has three custom-built core components: (1) an efficient real-time tracker that processes the videos from multiple cameras and detects the positions of the flies, (2) software that implements the stimulation rule, i.e., which positions or actions are to result in stimulation, and (3) a multi-channel LED controller to deliver light pulses according to the stimulation rule component’s instructions (Fig. 1a).

**Figure 1.**
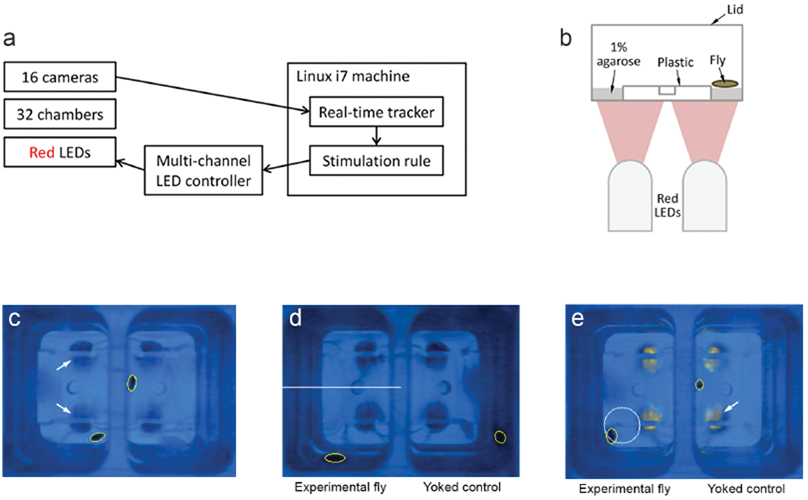
Overview of SkinnerTrax. **(a)** Overview of the SkinnerTrax components, with arrows indicating direction of information or control flow. **(b)** Schematic of the cross section of one chamber. We recorded through the lid with a camera above the chamber. **(c)** Sample frame from one of our videos. The frame shows two chambers, each holding one fly outlined by a yellow ellipse. Chambers are illuminated by white background light but appear blue due to filters placed in front of the camera (**Supplementary Fig. 1** and Online Methods). Arrows point to two of the red LEDs (in off state). **(d)** Sample frame during training in our differing-rate assay at a moment when the LEDs are off. The stimulation rate depends on whether the experimental fly is above or below the white line. The white line here and the white circle in the next panel were drawn by SkinnerTrax to visualize the stimulation rules for the users and were not visible to the flies. Note that we consistently paired each experimental fly with a so-called yoked control that receives stimulation simultaneously with the experimental fly, while when stimulation happens is controlled solely by the experimental fly’s behavior. Due to the simultaneous stimulation, a single channel (driving four LEDs) of our LED controller was sufficient per pair. **(e)** Sample frame during training in our enter circle assay with red LEDs in on state (arrow). The experimental fly just entered the white circle, causing stimulation of both flies.

Both the real-time tracker and stimulation rule components of SkinnerTrax are written in pure Python to make the code more accessible, while care was taken to perform all compute-intensive processing in fast libraries (primarily OpenCV) and to select efficient data structures and algorithms, allowing our Python code to run at essentially the speed of efficient native code. As a result, SkinnerTrax can easily handle 16 cameras at 320×240 pixels and 7.5 fps (our default) on a single Intel i7-4930K machine using only about 9% CPU. Compared to Ctrax^10^ 0.5.6, a popular non-real-time tracker which is also Python‐ and OpenCV-based, SkinnerTrax was about 8 times faster on one of our sample videos. (While designed for real-time tracking and capturing from cameras, SkinnerTrax can also be used on prerecorded videos.) Most of SkinnerTrax is operating system independent, but it was developed and is partly dependent on Linux. While Linux is not a real-time operating system, we analyzed several of our experimental videos, and in each and every case when the stimulation rule determined that the LEDs should be turned on, the next frame showed the LEDs to be on. For details on our multi-channel LED controller and additional details on the real-time tracker and stimulation rule components, see Online Methods. The code for all three components will be freely available at https://github.com/ulrichstern/SkinnerTrax.

In addition to the three core components, we used custom chambers to hold the flies. SkinnerTrax places few restrictions on the chamber design; in fact, the chambers we used were originally designed for high-throughput egg-laying experiments^11–13^ and worked well for SkinnerTrax, also. Each individual chamber held a single fly and was illuminated with two red LEDs for channelrhodopsin activation (Fig. 1b and **Supplementary Fig. 1**). (We generally used CsChrimson^1^ (CsC) for neuronal activation.) We captured two of the chambers per camera (Fig. 1c), for which we used inexpensive webcams. To simultaneously record many chambers, we used 16 webcams with each Intel i7 machine running SkinnerTrax (**Supplementary Fig. 2**, **Supplementary Video 1**, and Online Methods)

To illustrate that the stimulation rules in SkinnerTrax are highly flexible and can support a wide range of experimental designs, we next used it to implement three different behavior-dependent light stimulations. First, for our *differing-rate assay*, we made the stimulation dependent on the fly’s position. Specifically, the fly received red light pulses at different rates depending on where it was in the behavioral chamber (Fig. 1d). Second, for our *speed assay*, we made the stimulation dependent on the fly’s action. Specifically, the fly received red light continuously for as long as its speed of movement was below a user-defined threshold (**Supplementary Fig. 3a**). Third, for our *enter circle assay*, we made the stimulation dependent on both position and action. Specifically, the fly received a 250-ms pulse of red light whenever it performed the action of entering a circular area (i.e., crossed from outside to inside of the circle) (Fig. 1e). It is worth pointing out that since stimulation rules are implemented in software, once a SkinnerTrax setup is built, developing a new assay often requires no hardware development and is hence very fast.

Finally, we used our differing-rate assay to ask whether position-dependent stimulation of the bitter-sensing *Gr66a* neurons, whose activation has been shown to elicit avoidance behavior^6, 14^, can cause flies to exhibit active or memory-induced place avoidance. Using a 16-camera SkinnerTrax setup enabled us to examine 32 flies at a time. For details of the assay, see Online Methods. The experimental *Gr66a>CsC* flies showed clear positional preference for the less “punished” bottom half of the chamber during training sessions, while yoked controls showed no such preference (Fig. 2a–b and **Supplementary Fig. 3b**). In addition, the experimental flies showed positional preference for the bottom half during post-training breaks (Fig. 2b). We do not interpret the post-training preference as “learned place avoidance,” however, as the flies tended to move very little in the absence of stimulation and were much more likely on the bottom halves at the moments the training sessions ended. See Figure 2c for a typical trajectory of an experimental fly during training 1 and the post-training break. Interestingly, flies did increase their positional preference for the less punished half during training sessions as the number of session went up, suggesting their avoidance of light stimulation increased over time (Fig. 2d). Taken together, our results confirm that activation of Gr66a neurons elicits avoidance behavior and show that the avoidance behavior exhibits experience-dependent plasticity.

**Figure 2.**
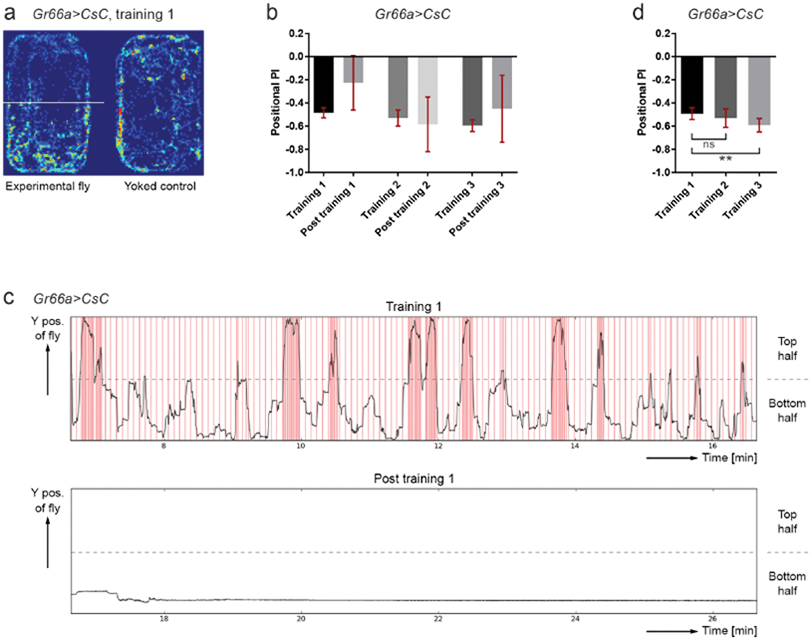
*Gr66a>CsC* flies in our differing-rate assay. **(a)** Sample positional heat map of one pair of *Gr66a>CsC* flies at the end of training session 1. For details of the assay, see Online Methods. Warmer colors indicate more time spent at the position. **(b)** Positional PI (preference index) of *Gr66a>CsC* flies. Positional PI = (NT - NB) / (NT + NB), with NT and NB denoting the numbers of frames the fly was in the top or bottom halves of the chamber, respectively. The bars show means with 95% confidence intervals. See also Statistics notes in Online Methods. N = 39 flies. **(c)** Sample trajectory of a *Gr66a>CsC* fly during training session 1 and the following post-training break. Red lines indicate red LED pulses, which occur at a higher frequency when the fly is in the top half of the chamber. **(d)** Positional PI of *Gr66a>CsC* flies during the last eight minutes of each ten-minute training session, with sessions excluded where the fly did not visit both chamber halves during the first two minutes of the session. The session exclusion combined with using only the last eight minutes of the session for the PI calculation avoids that the position of the fly at the moment the session started (which is non-random for later sessions) influences the calculated PI. Paired t-test, two-tailed, p = 0.34 (ns) and p = 0.0013 (**). The PI difference between trainings 1 and 3 passed normality. Using the Bonferroni correction to adjust for multiple comparisons does not change significance. N = 39, 39, and 38 sessions.

In conclusion, the efficiency of the SkinnerTrax code combined with the use of inexpensive components for video acquisition and light delivery makes achieving high throughput easy, speeding up experiments and even enabling previously impractical genetic screens. In addition, the flexibility SkinnerTrax provides for implementing stimulation rules allows for a wide range of different assays. The stimulation rule component currently has built-in support for recognizing only simple behaviors, but we plan to extend the range of behaviors by incorporating machine learning-based behavior recognition techniques similar to those that have recently been used for non-real-time video analysis^15, 16^.

## Methods

See Online Methods. (Below.)

## Acknowledgements

We thank the Duke Physics Shop for making our chambers, Texas Instruments for granting permission to copy a circuit diagram from their TLC59711 datasheet for Supplementary Figure 4b, and Kate Joy for comments on the paper. C.-H.Y. was supported by the National Institutes of Health under award number R01GM100027.

### Author Contributions

U.S. and C.-H.Y. conceived of the project, performed and analyzed the fly experiments, and designed stimulation rules. U.S. designed the software and hardware. C.-H.Y. designed the fly experiments. U.S. wrote the paper with help from C.-H.Y. All authors reviewed the manuscript.

### Competing Financial Interests

The authors declare no competing financial interests.

## Online Methods

**Camera-related hardware.** We used Microsoft LifeCam Cinema webcams and two Lagoon Blue (#172) filters from LEE Filters (Andover, UK) in front of each webcam to reduce red light intensity (**Supplementary Fig. 1**). We connected each set of four webcams to a plugable USB 2.0 4-Port Hub (model USB2-HUB4BC, Plugable Technologies, Redmond, WA). Note that a powered hub is required since four webcams typically require more power than what a single USB port can provide. Each hub was connected to our Linux i7 machine, an Intel i7-4930K machine put together from individual components and running Ubuntu 14.04. Since SkinnerTrax with 16 cameras (at 320×240 pixels and 7.5 fps) used only about 9% CPU on this machine, a much less powerful machine would likely have been sufficient.

For background lighting, we placed light pads (model LightPad 920, Artograph, Inc., Delano, MN) that were dimmed with 1.47 kΩ serial resistors to just 1% of their normal intensity under each apparatus (**Supplementary Fig. 1** and **2a**). We found the resulting white LED illumination of about 0.05 μW/mm^2^ in the chambers weak enough to not activate CsChrimson but strong enough as background light for tracking even with two Lagoon Blue filters. An alternative would have been to use infrared (IR) background lighting. In this case we would have had to remove the LifeCams’ factory-installed IR filters. Most non-night-vision webcams have such IR filters.

**Real-time tracker.** The tracking algorithm used is based on calculating the difference between the current frame and the background image – an estimate of the frame without flies – and using a detection threshold on the difference image. To calculate the background image in real-time, we used OpenCV’s accumulateWeighted() with some custom techniques to improve background image quality; e.g., we used a form of selective background update^17^ where pixels that are in the foreground according to a threshold less than the detection threshold are excluded from the background update, which reliably prevented resting flies from making it into the background. If more than one object that may be a fly is detected in a chamber (e.g., due to shadows), the tracker picks only the largest one and ignores the others, which relies on our “only one fly per chamber” setup and is similar to the shadow detection technique we used previously in non-real-time tracking^18^.

Despite the filters in front of the LifeCams to reduce red light intensity, the frames fell into two distinct lighting states depending on whether the red LEDs were on or off, which we previously addressed by calculating a separate background for each state^18^. For SkinnerTrax, we attempted to eliminate the need to calculate two separate backgrounds. We further reduced the difference between the two states by using only the blue channel of the video for tracking. To our initial surprise, however, the red LEDs typically appeared *darker* in the blue channel when they were on compared to when they were off, which was likely due to hue-preserving clipping by the webcam when the red or green channels would otherwise have exceeded their maximum value (255?) in areas of the frame showing the lit-up LEDs. By adjusting the saturation setting for each camera individually, we could make the red LEDs appear essentially identical in “on” and “off” states in the blue channel. (Adjusting, say, the brightness setting instead may have worked, also.) Combining blue channel tracking and saturation adjustment eliminated the need to calculate two separate backgrounds.

To make running experiments with many cameras easy, SkinnerTrax includes code that, for each camera when the experiment is started, automatically sets the camera settings (e.g., focal plane, exposure, and saturation) to saved good values and detects the precise position of the chambers in the video frame using template matching. The code to set the camera settings is LifeCam specific but should work with relatively minor changes for other webcams that are UVC (USB Video Class) compliant, which seems to typically be the case for recent webcams. For details of the template matching method, see our previous work in non-real-time tracking^18^.

**Stimulation rules.** Stimulation rules are implemented by directly writing Python code, which allows maximum flexibility in experimental design. SkinnerTrax uses threads and provides various utility functions to make implementing stimulation rules easier. Often an experiment consists of multiple sessions, each with an independent stimulation rule and with breaks in-between sessions, and one can readily implement the execution of a sequence of sessions (a “stimulation protocol”) in SkinnerTrax, fully automating the experiment. The execution of the stimulation protocol starts only once the fly has been detected by the real-time tracker, which requires some movement of the fly. If experimental flies are paired with yoked controls, protocol execution starts only once each fly has shown some movement.

**Multi-channel LED controller.** Our multi-channel LED controller (**Supplementary Fig. 2a** and **4**) is Arduino Uno-based and uses two Texas Instruments TLC59711 LED driver chips, providing a total of 24 independently controllable LED channels. It is worth noting that the number of channels can be increased readily by “chaining” additional TLC59711s. The TLC59711 uses pulse width modulation (PWM) to control LED intensity and an effective PWM frequency of about 19.5 kHz due to its “enhanced spectrum” feature. Note that the effective PWM period (~51 µs) was only a tiny fraction of CsChrimson’s off decay time (~20 ms)^1^, and the PWM should have no side effects in this application. For each channel used, we connected an LED cable with four Cree 624-nm 5-mm 15° LEDs (kit number C503B-RAN-CA0B0AA1) in series (**Supplementary Fig. 4a**). We used the lower Typical Application Circuit Example on page 2 of the TLC59711 datasheet with minimal changes for connecting the Arduino Uno and the TLC59711s (**Supplementary Fig. 4b
**). To simplify soldering, we used Adafruit’s TLC59711 boards (product id 1455, Adafruit Industries, New York City, NY). We also wrote a new Arduino library for controlling the TLC59711 (https://github.com/ulrichstern/Tlc59711) to fix problems with the library provided by Adafruit Industries.

On the Linux i7 machine, a single “LedController” process is in charge of communicating with the Arduino via serial port (over USB). We measured a roundtrip delay of about 4 ms, which matches the lowest numbers reported online. (If a reduction is required, a relatively simple switch from Arduino to Teensy would reduce the delay to about 1 ms.) Each of the real-time tracking processes communicates with the LedController process via TCP over a socket that is kept open throughout the run of the real-time tracker process to avoid connection establishment delays. The commands used are “set channel x to intensity y,” which work well for the continuous illumination (e.g., a 250-ms pulse) we used in our experiments. If pulse train illumination (e.g., ten 4-ms pulses at 40 Hz) is required, the Arduino code could be extended to support pulse trains.

**Handling many webcams.** The Microsoft LifeCam Cinema reserves about 48% of USB 2.0 bandwidth independent of the resolution and frame rate used, which typically limits the number of LifeCams to just one or two per USB controller. The problem is not limited to the LifeCam Cinema or Microsoft; several other webcams also reserve more bandwidth than they need. In theory, for the resolution and frame rate we used (320×240 pixels, 3 bytes per pixel, 7.5 fps), the USB 2.0 bandwidth of 480 Mbit/s allows up to 34 webcams. Fortunately, the Linux UVC driver includes a so-called “quirk” to ignore the bandwidth reservation from the camera (UVC_QUIRK_FIX_BANDWIDTH), allowing more than two LifeCams to be used per USB controller (http://stackoverflow.com/q/25619309/1628638). Even using the quirk, the actual number of supported cameras per USB controller can still be much smaller than the theoretical maximum, and adding additional USB controllers via USB cards can also increase the number of cameras supported; we added one ANKER USB 3.0 card (model 68UPPCIE-4SU, Anker Technology Co., Limited, Hong Kong; we had good experiences with the VIA VL805 USB controller it uses) to our i7 machine.

In addition, current versions of OpenCV (both 2.4.13 and 3.1.0) by default support at most eight webcams on Linux. But this can be increased easily by changing constant MAX_CAMERAS in the OpenCV source code and rebuilding OpenCV (http://stackoverflow.com/q/38619801/1628638).

Note that the requirement to handle many webcams was partly due to the high sidewalls of our chambers. With low-sidewall chambers, the number of chambers per camera could be increased, reducing the number of cameras required for high throughput.

**Differing-rate assay.** For our differing-rate assay, we programmed SkinnerTrax to administer three 10-min training sessions, each followed by a 10-min “break” without light pulses, and an initial 5-min break before the first session. During training sessions, flies received 250-ms red light pulses of about 11 μW/mm^2^ (25% LED intensity) at a rate of 1 Hz when they were in the top half of the chamber and at a lower rate of 0.2 Hz when they were in the bottom half (Fig. 1d and 2c). We chose a low stimulation rate of 0.2 Hz over “no stimulation” for the bottom half since *Gr66a>CsC* flies tended to move very little during periods without stimulation that followed a period where they had been stimulated multiple times.

**Statistics notes.** We used GraphPad Prism 6 to perform statistical tests and significance level α = 0.05. In the figures, bars show mean PIs (see legend for Fig. 2b) with 95% confidence intervals, which can be used to assess how different the mean PIs are from zero (no preference). We used ** for p < 0.01 and “ns” for not significant.

